# Eye-tracking for low vision with virtual reality (VR): testing status quo usability of the HTC Vive Pro Eye

**DOI:** 10.1101/2020.07.29.220889

**Authors:** Alexandra Sipatchin, Siegfried Wahl, Katharina Rifai

**Author notes:** Corresponding author: Alexandra Sipatchin, ZEISS Vision Science Lab, Universitätsklinikum Tübingen, Elfriede-Aulhorn-Str. 7, 72076 Tübingen.

## Abstract

**Background:** Adding an eye tracker inside a head-mounted display (HMD) can offer a variety of novel functions in virtual reality (VR). Promising results point towards its usability as a flexible and interactive tool for low vision assessments and research of low vision functional impairment. Visual field (VF) perimetry performed using VR methodologies evidenced a correlation between the reliability of visual field testing in VR and the Humphrey test. The simulation of visual loss in VR is a powerful method used to investigate the impact and the adaptation to visual diseases. The present study presents a preliminary assessment of the HTC Vive Pro Eye for its potential use for these applications.

**Methods:** We investigated data quality over a wide visual field and tested the effect of head motion. An objective direct end-to-end temporal precision test simulated two different scenarios: the appearance of a pupil inside the eye tracker and a shift in pupil position, known as artificial saccade generator. The technique is low-cost thanks to a Raspberry Pi system and automatic.

**Results:** The target position on the screen and the head movement limit the HTC Vive Pro Eye’s usability. All the simulated scenarios showed a system’s latency of 58.1 milliseconds (ms).

**Conclusion:** These results point towards limitations and improvements of the HTC Vive Pro Eye’s status quo for visual loss simulation scenarios and visual perimetry testing.

## Introduction

Eye-tracking is a recognized technique used to investigate the relation between eye movements and human cognition. Its first use dates back to the 19th century when Huey (1898) and Delabarre (1898) for the first time traced eye movements on a rotating drum. Buswell (1935) reported the first systematic exploration of the fixation position. In comparison, eye tracking in VR is a relatively new field with its first application in gaze-contingent studies, starting only two decades ago (Danforth et al., 2000; Tanriverdi and Jacob, 2000).

Gaze-contingent studies use actively or passively the gaze as an input. For gaze-contingent active scenarios, gaze can actively perform an action and substitute a mouse to select stimuli, open menus. Passive contingent studies use gaze to change a display dynamically. For example, for foveated rendering, the central part of the screen maintains a high resolution. In contrast, the peripheral part is being updated with a lower amount of detail before the end of each change in gaze position (Holmqvist et al., 2011). Most of the studies assume that the participant is not aware of the changing display since during a saccade stimuli are invisible (Brooks and Fuchs, 1975; Riggs et al., 1982). Human saccadic movements appear to be stereotyped (Becker, 1989) and accurate, even beyond 60 years old (Warabi et al., 1984; Munoz et al., 1998). Saccades are very brief eye movements, and their duration depends upon the amplitude. For reading, saccades last from 20 to 40 milliseconds (ms) (Rayner et al., 2001) and a 5°, 10°, 15°, and 20° saccade have a mean duration between 31 and 54 ms (Baloh et al., 1975; Bahill et al., 1981; Becker, 1989; Thickbroom et al., 1991; Behrens et al., 2010; Gibaldi et al., 2017).

For active gaze-contingent paradigms, eye-tracking accuracy and precision must be high so that the data samples are being correctly identified as belonging to the displayed button area for as long as it is being fixated (Holmqvist et al., 2011). Accuracy is the average angular error between the measured and the location of the intended fixation target. Precision is the spread of the gaze positions when a user is fixating a known location in space (Holmqvist et al., 2011; Gibaldi et al., 2017). For passive applications, the system’s latency has to be lower than the saccade duration. Temporal precision is the average end-to-end delay from the tracked eye’s actual movement until the recording device signals that the movement has occurred (Holmqvist et al., 2011). Passive gaze-contingent scenarios need an eye tracker with good temporal precision.

Research about eye-tracking data quality in VR is limited (Lohr et al., 2019), with most studies investigating the head-mounted display’s (HMD) tracking accuracy (Niehorster et al., 2017; Borges et al., 2018; Peer et al., 2018; Groves et al., 2019). The present study represents a pilot study to evaluate the usability of the HTC Vive Pro Eye integrated eye-tracker (Vive, 2019a) for a visual field (VF) testing and simulation of low vision.

According to the International Classification of Disease, 10th Revision (ICD-10) low vision coincides with a visual acuity value of less than 6/13 (0.3) but equal or higher than 3/60(0.05) for the better eye using correction (Thylefors et al., 1995). Disorders that cause low vision are age-related macular degeneration (AMD), glaucoma or retinitis pigmentosa (Donders, 1957; Berger and Porell, 2008; Jager et al., 2008; Mantravadi and Vadhar, 2015). Symptoms include blurred vision, central and/or peripheral VF loss. A VF test can identify damages in central and peripheral vision covering a visual field up to +-30° temporally and nasally.

The concept of VR for VF perimetry testing appeared in 1998 (Kasha, 1998). Since then, studies tested its comparability to the standard Humphrey visual field analyzer (HFA, Carl Zeiss Meditec Inc, California, USA) with mixed results (Hollander et al., 2000; Plummer et al., 2000; Tsapakis et al., 2017; Erichev et al., 2018; Mees et al., 2020). Gaze tracking for VR VF testing adds new resources for the advancement of mobile and portable perimetry. Eye-tracking makes it so that stimuli can appear at the fixation area, and it can implement more objective criteria to capture responses similar to active gaze-contingent paradigms, like, for example, a visual grasp mode (Wroblewski et al., 2014). Visual grasp is based on the so-called eye movement perimetry. It overcomes long periods of fixation of peripheral stimuli common to standard perimetry (Trope et al., 1989). During central fixation, a stimulus appears and induces an automatic reflex towards the new target. When the gaze change is consistent with the new target position, the test identifies that part of the visual field as intact, and it moves forward. After correct identification, the new stimulus becomes the new fixation. This is the idea behind a hand-free visual grasp mode where the gaze replaces the patient’s response (Wroblewski et al., 2014).

Visual impairments simulations in VR safely investigate the effect and the adaptability to visual defects. Simulations of the appearance of the visual loss allow controlled and comparable effects across subjects (Bertera, 1988; Bertera 1992). These studies use passive gaze-contingent applications that take the gaze input to simulate low vision on a display that continuously updates where the subject is looking. Extensive studies showed that normal subjects develop similar visual strategies to adapt to low vision as patients do (Bertera 1992; Zangemeister and Oechsner, 1999; Kwon et al., 2013; Walsh and Liu, 2014; Geringswald and Pollmann, 2015; Barraza-Bernal et al., 2017a; Barraza-Bernal et al., 2017b) and can be used to study their impact (Barraza-Bernal et al., 2018).

A head-still condition investigates the HTC Vive Pro Eye for a VF VR usability. The test examines data quality across a wide visual field to check for limits from target position on screen (Feit et al., 2017). A head-moving situation tests the system under free head movement to investigate possible low precision and/or data loss induced by movement (Holmqvist et al., 2011; Holmqvist et al., 2012; Niehorster et al., 2020).

For its applicability for visual loss simulations, a direct, objective, and automatized end-to-end method examines the temporal precision, based on existing end-to-end tests (Reingold, 2014; Saunders and Woods, 2014; Gibaldi et al., 2017). Direct end-to-end measurements of latency are more reliable compared to non-direct tests since interactions between different systems are hard to predict (Saunders and Woods, 2014). An artificial pupil and a saccade generator are used to examine the average time between the onset of an artificial eye and successive saccade and the display of the first eye-tracking data showing a change on the display. Unlike the use of a human observer, the major advantages are identical motion sequences that can be reproduced multiple times and tight control over the input given to the tracker without additional delays that can be induced by a subjective input, such as a human observer (Reingold, 2014).

## Materials and methods

### Participants

Eleven participants took part in the data quality assessment test (6 females and 5 males, mean age 28.73, standard deviation (SD): ±2.49 years, 5 with bright eye color and 6 with dark; 6 having previous experience with the eye tracker; 8 wearing eye-correction, 3 wearing contacts and 5 wearing corrective glasses, the rest did not need correction). The direct end-to-end method for latency required no participants.

### HMD: eye-tracking and display

We collected eye-tracking data in VR using the built-in Tobii eye tracker (Core SW 2.16.4.67) with an autonomous eye-tracking algorithm processing integrated (Tech, 2019) and a sampling frequency of 120 Hz (Vive, 2019a).

Nine near-infrared light (NIR) illuminators per eye illuminate the eye for pupil center corneal reflection (PCCR) for non-intrusive eye tracking (Pro, 2015a). One infrared (IR) camera per each eye, placed inside the lens-display tube, captures the reflections (Wiltz, 2019). The image captured is used to identify the provoked reflection patterns on the cornea and pupil and calculate a vector between the two reflections. The direction of this vector represents the gaze direction. The illumination technique used is a proprietary illumination method built to collect eye-tracking data from a population wearing glasses, hard and soft contact lenses. Tobii Pro SDK v1.7.1.1081 (Pro, 2015b) and Vive SRanipal SDK v1.1.0.1 (Vive, 2019b) are used to access non-filtered and filtered eye-tracking data, respectively. The system’s accuracy estimation is 0.5° to 1.1° within a field of view of 20° (Vive, 2019a).

The HMD has two AMOLED screens, with a resolution of 2.880 × 1.600 pixels in total (1.440 × 1.600 pixels to each eye resulting in a pixel density of 615 pixels per inch (PPI)), with a refresh rate of 90 Hz and a field of view of 110° (Vive, 2019a).

### Calibration procedure

HTC VIVE Pro Eye uses the Super Reality (SR) runtime to enable eye tracking. It offers calibration with five points. Calibration starts with a central point that shrinks to focus the gaze. It moves the other four points from one point to the next, shrinking and then disappearing to the next position. The eye tracker waits until data is collected. Calibration was carried out successfully for all participants.

### Set-up

For the virtual experiment, we used the Unity 2019.1.10f1 version as a design tool, with C# as a programming language, running on a PC with Windows 10 Home, having a 64-bit operating system, an Intel Core i7 −7700HQ, 2.8 GHz, 16 GB RAM, and a NVIDIA GeForce GTX 1070 GDDR5 graphics card. We used a single-board computer, the Raspberry Pi (Raspberry Pi, model B 2018, full-price under 50 euros) controlling a Raspberry Pi camera (Version 2.1, with the capability of 120 Hz, full-price under 30 euros) for the end-to-end direct latency tests.

### Experimental procedure

#### Data quality measurements – Head Still and Head Free tests

We tested accuracy and precision in a virtual environment where fixation targets (Figure 1A) were two concentric circles, one internal black and one external red circle with a radius of 0.72 degrees of visual angle, positioned at 1 meter in a Unity world coordinate system. The target would appear at 25 different sample positions distributed across 5 columns and 5 rows covering a visual field of °±26.6 (Figure 1B). We investigated two separate conditions: head-still and head-free. In the first condition, the target position was fixed to the HMD, and subjects had to keep their head still, saccade to an appearing target, and fixate it. In the head-free condition, targets were positioned in a world-fixed coordinate system, and we instructed subjects to saccade towards the appearing target, fixate it and then move their head naturally, while fixating, towards the position where it appeared. Subjects performed the task in both conditions in a seated position on a chair. We randomized the target position, and we displayed the target for 5 seconds (Clemotte et al., 2014) with 5 repetitions (5 sec/target*25 targets*5 repetitions= 625 seconds, approximately 10 min and a half). In the head free condition, we added a central fixation target (coordinates: [0,0,0]) at the end of each target presentation that lasted 2 seconds. Central fixation after each fixation was added to make the participates come back to the same referencing point (5 sec/target+2 sec/central target*25 targets*5 repetitions = 875 seconds, approximately 15 minutes). We used the Tobii Pro SDK to access non filtered data to avoid alteration of eye-tracking samples (Holmqvist et al., 2017; Orquin and Holmqvist, 2018).

**Figure 1:**
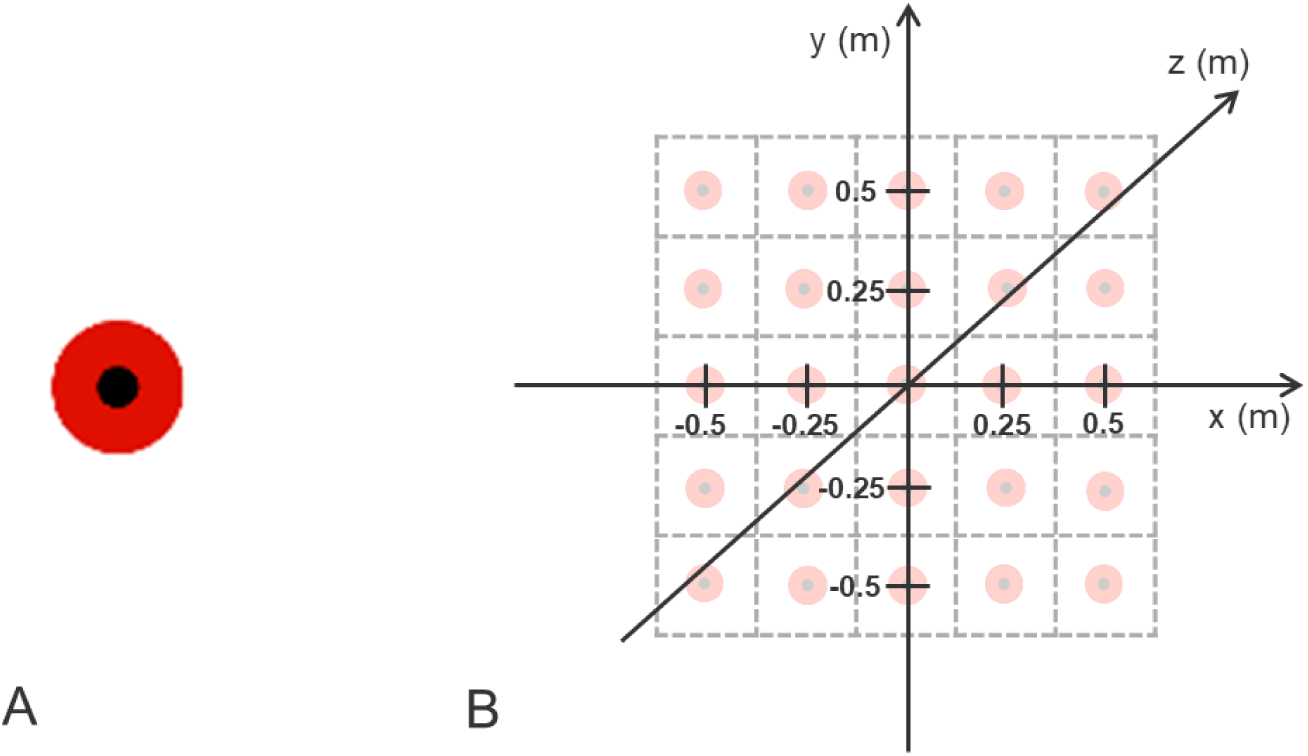
Schematic representation of the data quality assessment set-up. The target (A) is a virtual object with two concentric circles, one internal black one and one external red. Targets are distributed across 5 rows and 5 columns (3D wold coordinate system, B) with origin the centre of the HMD.

#### Temporal precision measurements – Eye-detection and Gaze-contingent tests

The method uses a low-cost configuration (Figure 2A): a Raspberry Pi single-board computer controls the output of IR light-emitting diodes (LED) and records, with a Raspberry Pi camera, the resulting eye-tracking events displayed by the HMD.

**Figure 2:**
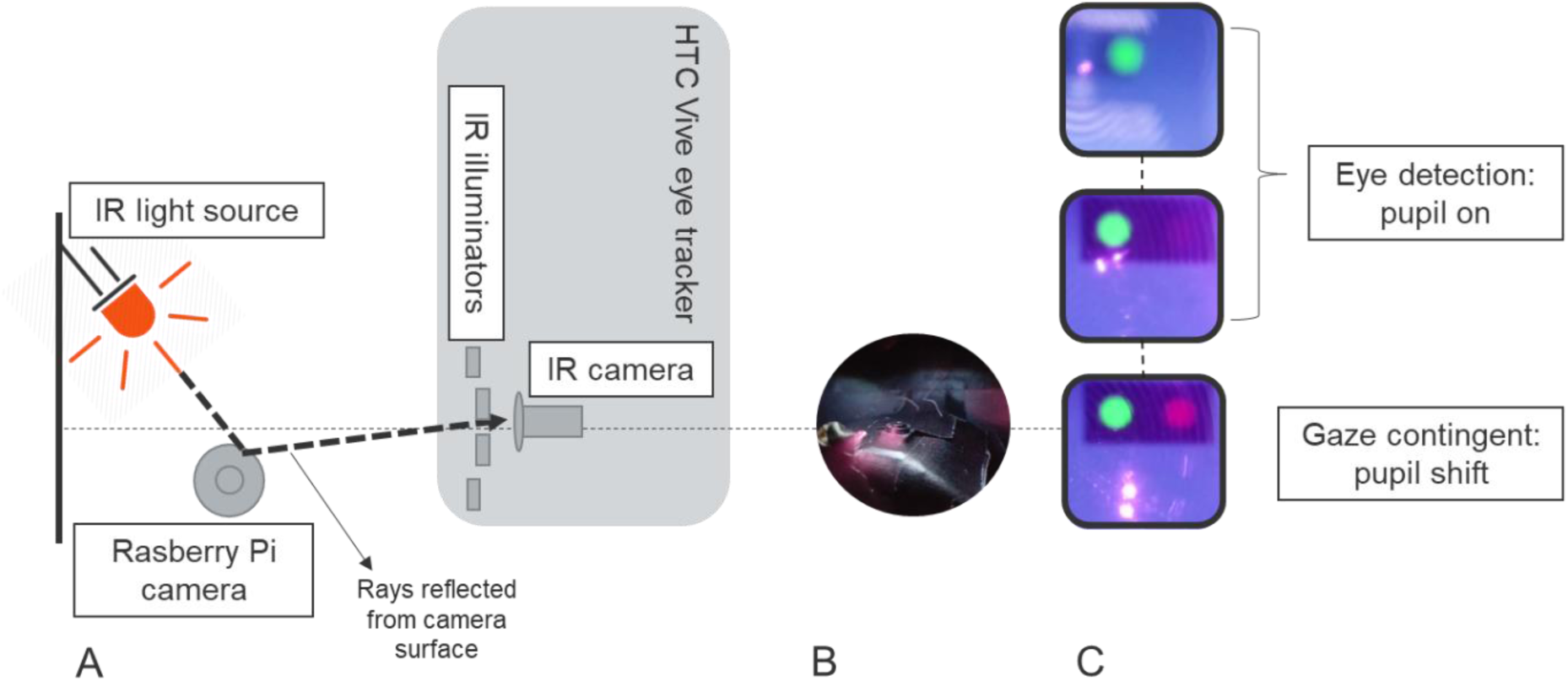
Schematic representation of the temporal precision set-up. IR light source (orange) represents the IR LEDs illuminating the Raspberry Pi camera (A, schematic and B, picture of the camera). The IR camera inside the HTC Vive Pro Eye captures the reflected rays by the Raspberry Pi camera (dotted lines). The Raspberry Pi camera can record the reflection as two separate events: as a reflection on the HMD lenses (pink dot), and as an artificial eye, the colored big dots (C). Upper and middle images are the eye-detection scenario, the last one is the gaze-contingent one. We collected data using the Tobii Pro SDK for the upper one and the SRanipal SDK for the last two ones.

**Figure 3:**
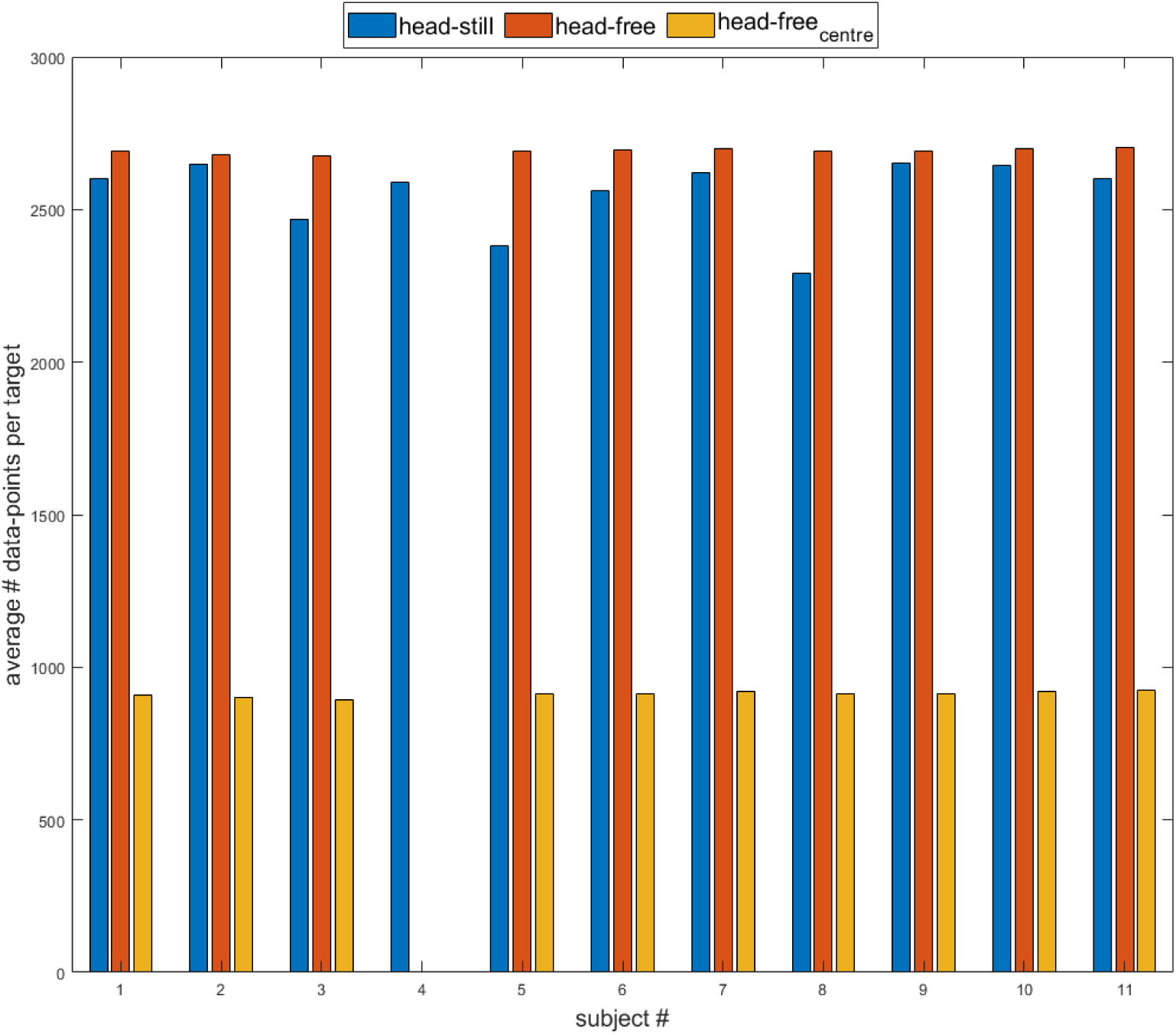
Bar Plot for the average number of data points per target. Blue bars represent data for the head-still condition, and the orange and yellow bars are the data for the head-free one. The orange ones are the data collected for the 25 targets, and the yellow bars are the points collected for the central referencing target.

The method tricks the eye tracker into two different scenarios: first into the detection of an eye, the eye-detection scenario, then next into an abrupt change in gaze position of the recognized artificial pupil, the gaze-contingent scenario. The first part of the study checks differences in latency when identifying an appearing eye between two different SDKs: Tobii Pro SDK v1.7.1.1081 and Vive SRanipal SDK v1.1.0.1 (Pro, 2015b; Vive, 2019b). The second scenario introduces a modified version of an artificial saccade generator (Reingold, 2014).

We used a virtual environment running on the PC which displayed ongoing successful and correct eye tracking. We used the Raspberry Pi for the eye-detection scenario to turn-on two IR LEDs and illuminate the Raspberry Pi camera for 1 second. This lead to clear IR reflections from the camera (figure 2A and B) and the HMD (Figure 2C). The reflections from the camera lead to a pupil-on event: the appearance of a green dot (Figure 2C). In this scenario, Tobii Pro SDK and SRanipal were both used to display the pupil-on event. We used the VR Positioning Guide Prefab, incorporated in the Tobii Pro SDK (Figure 2C, upper image) and a similar programmed version of the Prefab for the SRanipal SDK (Figure 2C, middle image).

For the gaze-contingent scenario, we used only the SRanipal SDK to display the pupil-on event and generate an artificial saccade. We placed two additional IR LEDs at a 1cm distance from the other two. The second pair of IR LEDs simulated an abrupt change in gaze position of the previously recognized artificial pupil. The Raspberry Pi turned off the first two at the same time. This produced the pupil shift event: an appearance of a bright red dot (Figure 2C, lower image). The pupil shift event did not disrupt the first pupil-on event since the display of this event was programmed such that a green dot should still be shown as long as an eye is being detected.

The Raspberry Pi camera controlled through the Raspberry single-board computer recorded the events displayed by the HMD (Figure 2C).

We recorded ten different videos for both scenarios, each with warming up camera period of 1 sec and an interval of 2 seconds turn off of all LEDs. The Raspberry Pi produced 33 IR LED on-off trials for each video when using the Tobii Pro SDK leading to 330 repetitions and a recording time of 16.7 minutes. For the SRanipal SDK, 660 repetitions were recorded with a total time of 21.7 minutes with each scenario having 33 IR LED on-off.

## Data analysis

### Data Pre-Processing: Eye-tracking accuracy and precision

The data provided by the Tobii Pro SDK Save Data Prefab at each sample data are HMD position and rotation, HMD-local eye position (vector of eye position measured in millimeters from the center of the HMD) and HMD-local gaze direction (a normalized vector re-referenced in HMD’s coordinate system pointing from the pupil towards the virtual object) both for the left and right eye separately. It also provides additional gaze origin and direction vectors to indicate pupil position and direction transported into world coordinates. HMD’s position vector and rotation quaternion are used to recalculate eye’s position (rotated HMD-local eye position around the rotation quaternion is added to HMD position), while HMD-local gaze position is used to re-reference the new gaze direction (similar calculation to Clemotte et al., 2014, Eye-Gaze, normalized vector in this case).

For each sample, we averaged each HMD’s local gaze coordinates for direction and origin. The same calculation is performed for the world-eye data automatically by the Prefab. In the head-still condition, we used the average of the HMD-local gaze direction and position vector, while in the head-free condition we took the world gaze already averaged vector as provided by the Tobii Pro SDK Prefab. The target position was saved at each sample, as defined by the experimental procedure in 3D Unity coordinates (Figure 1B). For each condition, the targets were then re-referenced to the eye by subtracting the eye’s position vector from the target’s coordinates (target vector - eye position vector).

For each sample data, we calculated the angle between the gaze direction vector (GD) and target-eye vector(TE) using the same formula (1) as Clemotte et al., 2014 to estimate the angle between two vectors (angleV).

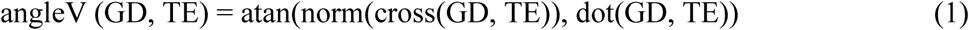

Atan calculates the inverse tangent, norm normalizes the vector, cross, and dot calculate the cross and the dot product respectively.

In the head-free condition, for each data sample, to separate between fixation during head-non-moving (Free__stable_) and fixation during head-moving phases (Free__moving_), we took the differential of the speed of the HMD’s rotation quaternion rotated around a normalized vector. For the analysis, we discarded the first 500 ms (Clemotte et al., 2014) after the target appearance that was considered as the time a subject used to direct the gaze towards it. In both conditions, we excluded from the analysis gaze points where no eye could be tracked both for the left and the right eye before averaging to avoid large errors in the mean’s calculation. For the data loss analysis, we kept gaze points where no eye was detected.

### Data analysis: Eye-tracking accuracy and precision

Accuracy is defined as the mean of all the angles (angleV) calculated between GD and TE using the formula described in (1). We used the common practice to calculate the spatial precision of the eye-tracker (Blignaut and Beelders, 2012), the root mean square (RMS) of the inter-sample angular distances between successive GDs. As an additional precision indicator, we also plotted a bivariate contour ellipse area (BCEA) for left, right, and the average of the two eyes to show the area that encompasses 50% of fixation points around the mean for each given target.

For the head-still condition before averaging between the two eyes, we conducted a one-way ANOVA test to check for differences in accuracy between the two eyes. We computed an overall average and an average for different percentiles of users for accuracy and precision. We calculated percentiles to analyze changes in eye data quality across the population tested.

A one-way ANOVA way tested how eye tracking data differs across screen regions with the horizontal line as the independent factor and the vertical ones as levels. Differences observed across the horizontal line might be an indication of the altering of eye-tracking data quality induced by reflections from vision corrections (Dahlberg, 2010).

In the head-free condition, we calculated the average precision as RMS and a one-way ANOVA tested how precision is affected by phases of stable head and moving head while subjects fixated the target. We also estimated the amount of data loss during the two phases. The data loss percentage was calculated using a similar formula (2) as Niehorster et al., 2020:

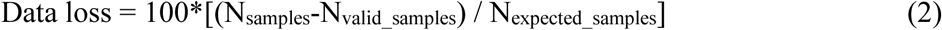

where N_samples_ represent the number of data samples recorded after the exclusion of the initial 500 ms and N_valid_samples_ are the number of samples during which a valid gaze position was recorded.

### Data analysis: Temporal precision

We converted the recorded videos into images frame-by-frame through a converter program (Free Video to JPG Converter, version 5.0.101). We programmed an automatized method to detect the elapsed frames between the onset of the LEDs and the onset of the different dots. We used the Color Thresholder app from the Matlab Image Processing Toolbox (version 10.4) to manipulate the color components of sample frames via a hue, saturation, value (2HSV) color space. We created three separate RGB 2HSV segmentation masks: one for the LED’s reflection on the HMD, one for the appearance of the green dot, and one for the appearance of the bright red dot (Figure 4). The masks indicated how many pixels in the frame contained the events. We created a script to count the number of frames between LEDs and the green dot onset and LEDs and bright red dot onset. For each frame, whenever the green and bright red dot were on or when the LEDs were on while using the Tobii Pro SDK, the script attributed a flag for a number of pixels greater than 10. When using the SRanipal, to differentiate between the first and the second pair of LEDs on, for each frame, the script attributed a flag whenever the number of pixels was greater or smaller than given values. This was possible since the second pair of LEDs cause a bigger reflection area (Figure 4, 2^nd^ LED). The script identified the first LED pair when the number of pixels was greater than 10 and smaller than 250. A number greater than 300 indicated the second pair of LEDs on. For the eye-detection scenario, both when using the Tobii Pro SDK and the SRanipal SDK, the script counted the flags from the LEDs onset until the green dot onset. For the gaze-contingent scenario, the count started with the second pair of LEDs onset and ended with the bright red dot appearance.

**Figure 4:**
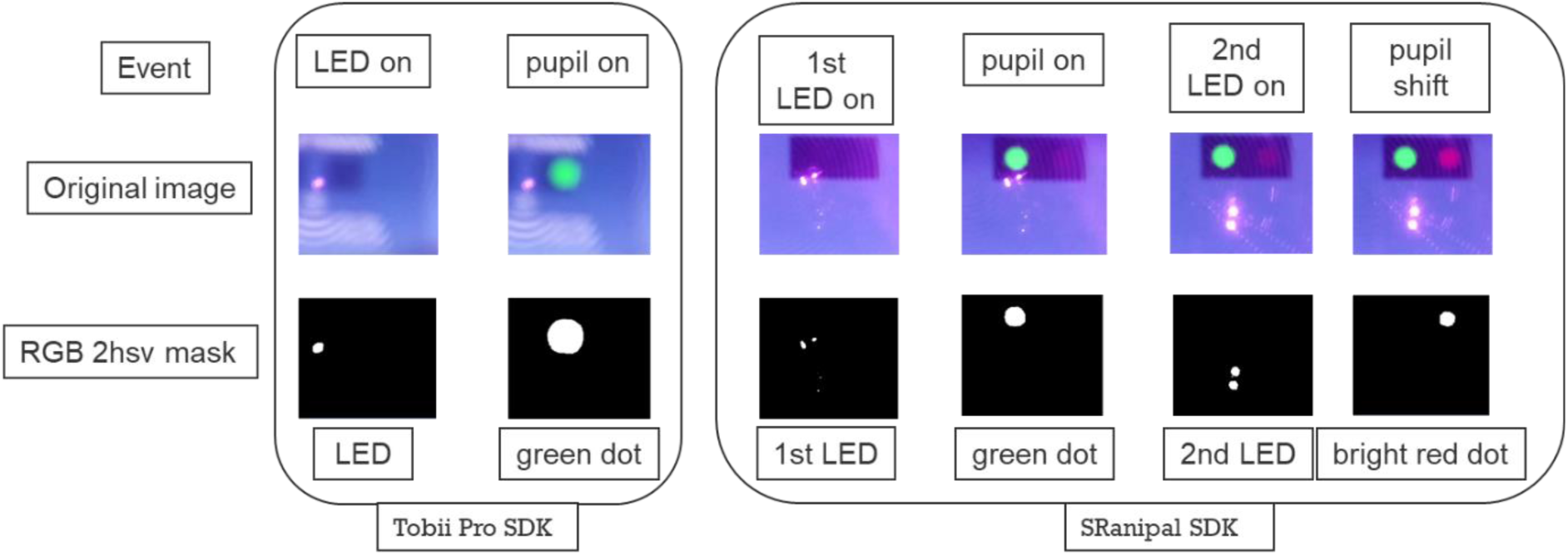
Matlab generated a segmentation mask (RGB 2HSV) for events selection. Upper images, frames containing the different events. Lower images: output from Matlab’s image segmentation mask (RGB 2HSV) of the selected events. The masks automatically identified the LED and dot events.

We plotted a histogram with the resulting intervals between events and tested for normal distribution with a one-sample Kolmogorov-Smirnov test. A boxplot is also plotted to compare the different scenarios displayed through the two SDKs. Temporal precision is calculated as the median of frame numbers elapsed between the LED and the different dot event multiplied by the mean duration of each video frame.

## Results

### Results: Pre-Processing Eye-tracking accuracy and precision

After data selection for each target, subjects had an average of 2550 data points in the head-still condition and 2692 points in the head-free condition; the central fixation target, used as a referencing point in the head-free condition, had 912 points (Figure 3).

### Results: Eye-tracking accuracy and precision

In the head-still condition, the one-way ANOVA resulted in no significant differences in accuracy between the two eyes (F (1,20) = 0.81, p = 0.38; mean left eye: 4.16° SD: ±1.49 and mean right eye: 4.75° SD: ±1.63; Figure 5 and 6). For this reason, we used the average across eyes (mean average both eyes: 4.16°, SD: ±1.40) for the analysis. Precision has a mean of 2.17°, SD: ±0.75. The BCEA shows that the accuracy and precision of the estimated gaze are worse at the most outer horizontal regions and that the central line has higher accuracy and precision than the most externally positioned targets, with the highest level of accuracy and precision for the central target (Figure 7). Comparing horizontal regions, a one-way ANOVA revealed that there is a significant difference in accuracy and precision (F (4,50) = 3.35, p = 0.02 for accuracy; F (4,50) = 3.6, p = 0.01 for precision). Post-hoc t-tests (Bonferroni corrected) show the center as being more accurate than the upper horizontal (p<0.03, central row mean offset: 2.26°, SD: ±0.73; upper row mean offset: 6.16° SD: ±5.50), and as more precise than the lower horizontal (p<0.01, central row RMS mean: 1.63° SD: ±0.30 and the lowest row RMS mean: 3.15°, SD: ±2.00). We plotted fixational eye movements and subjective data revealed unstable fixation patterns for the upper row (Figure 8) and deviations for the lower one (Figure 9).

**Figure 5:**
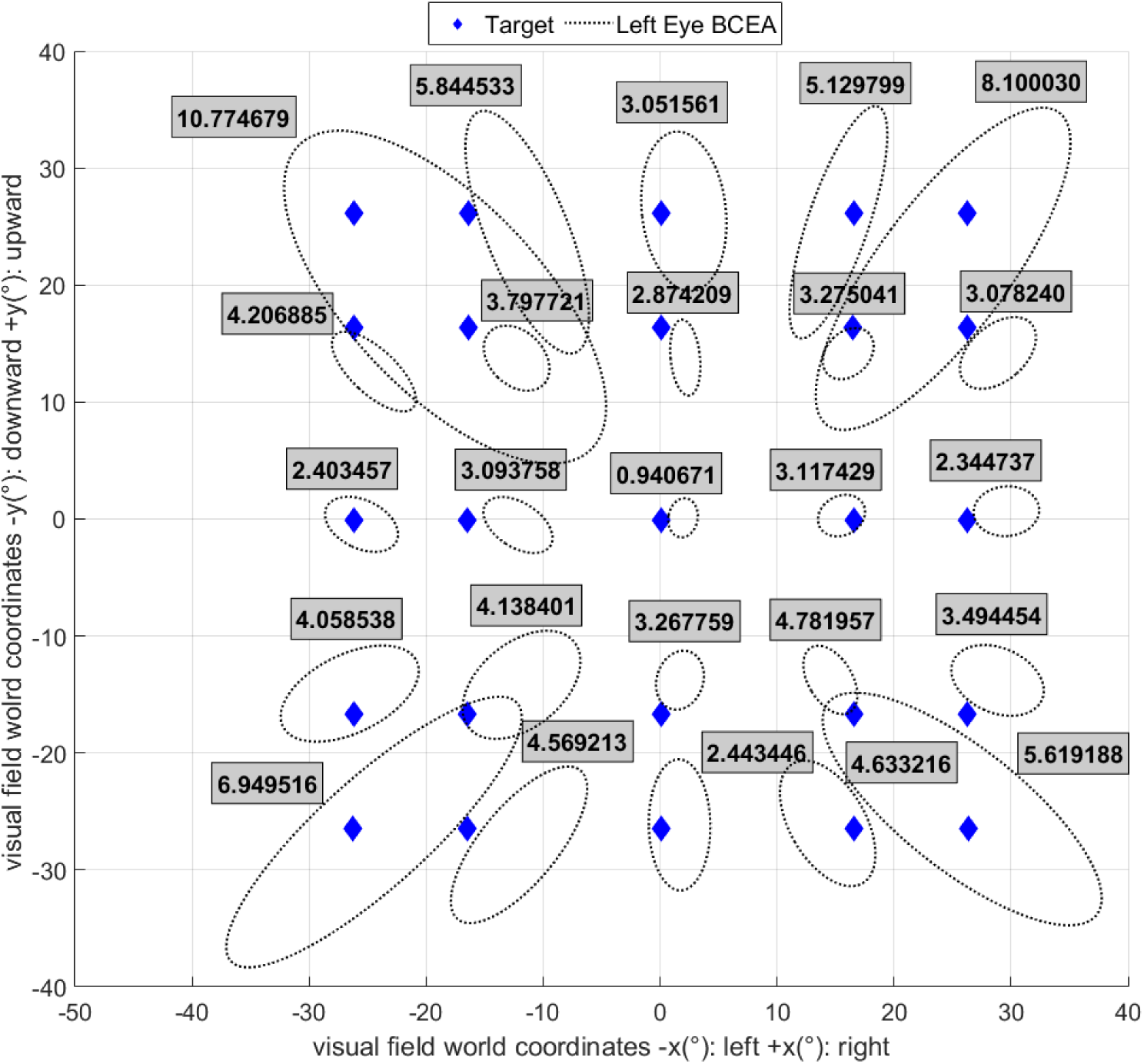
BCEA of the estimated gaze points for the left eye. Covariance ellipses (dotted lines) are fitted to the left eye’s gaze points corresponding to all fixations of the same target across all subjects. The values inside the grey box are the left eye’s average offset per target. The blue diamonds are the targets.

**Figure 6:**
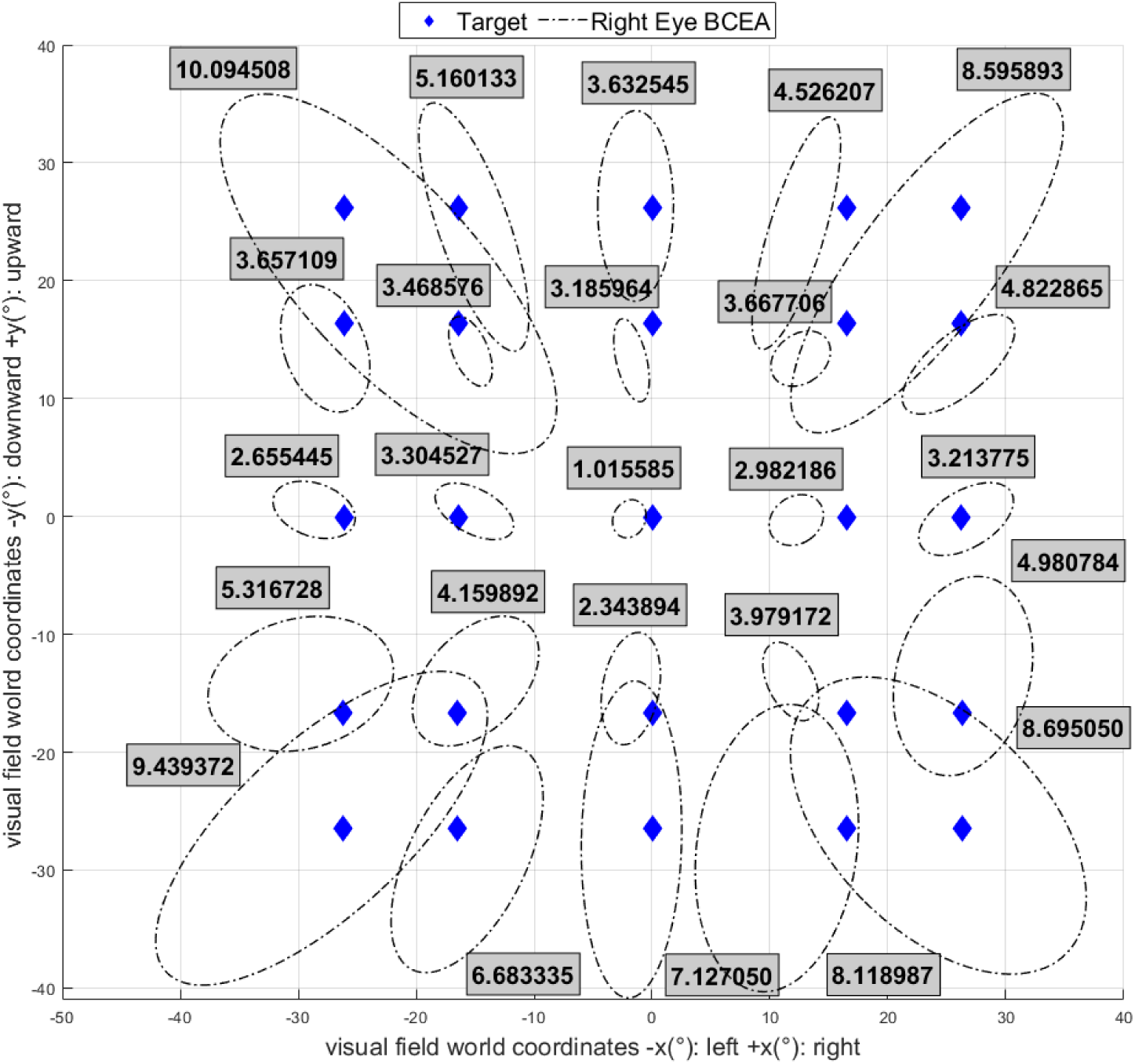
BCEA of the estimated gaze points for the right eye. All fixations for the right eye of the same target (blue diamond) across all subjects are fitted to covariance ellipses (dotted lines) and the grey boxes contain the average offset value per each target.

**Figure 7:**
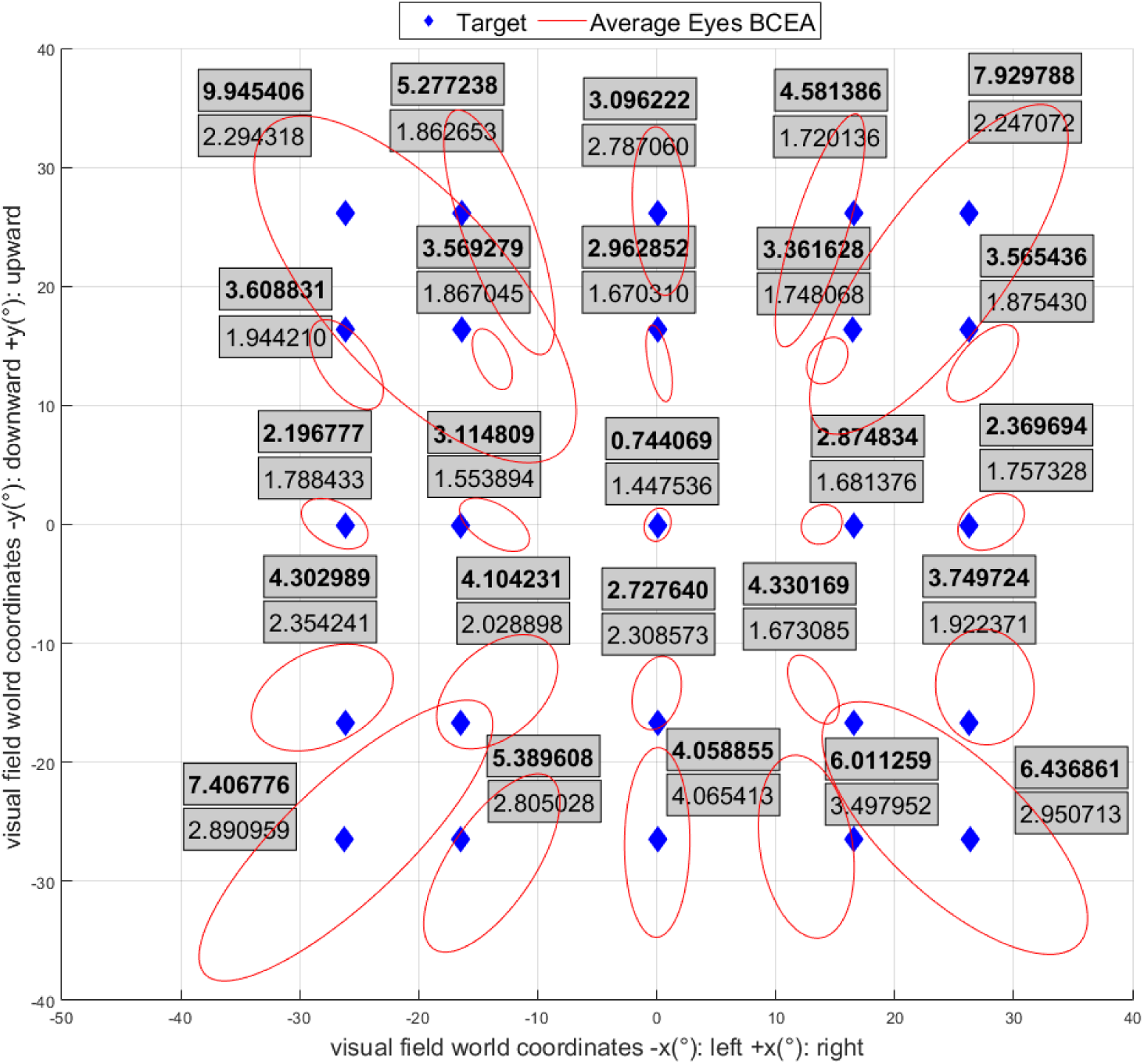
BCEA of the estimated gaze points for the average of both eyes. Covariance ellipses of the average of both eyes (red lines) are fitted to the gaze points corresponding to all fixations of the same target across all subjects. The grey box shows mean offset (in bold) and RMS (normal) for each target.

**Figure 8:**
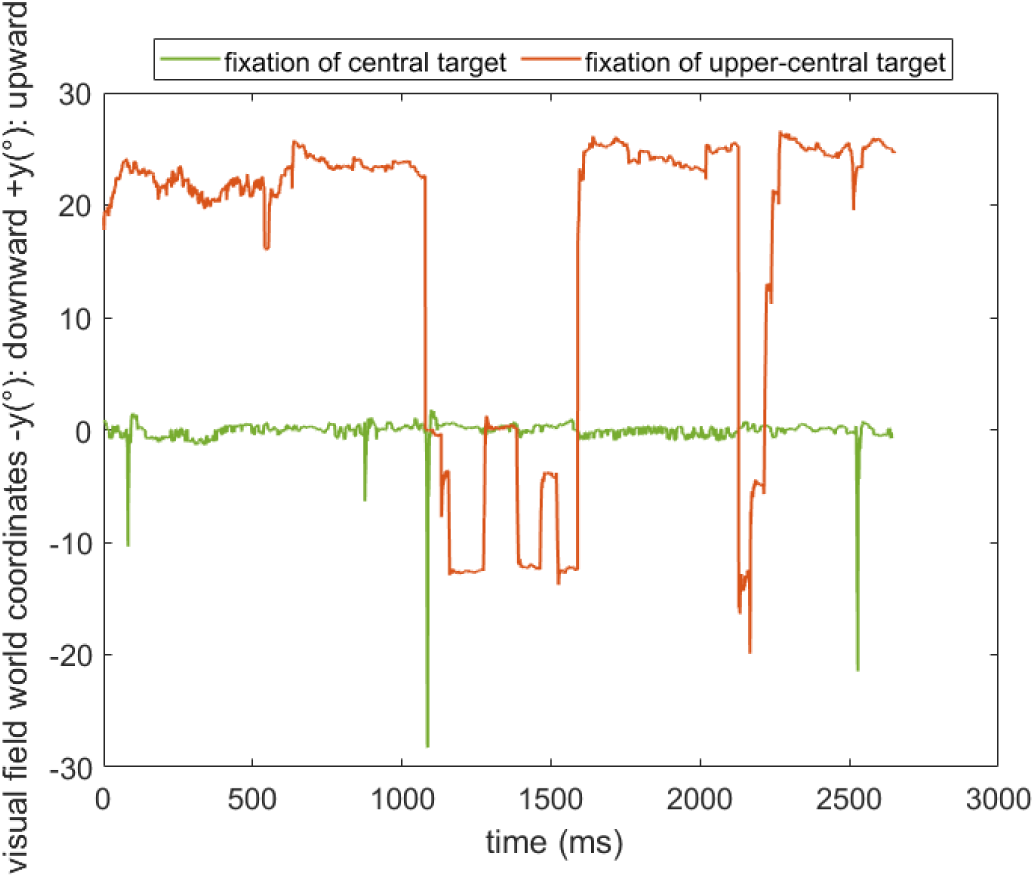
Subjective fixational data of targets positioned in the first row. Fixations plotted across time evidence an unstable fixation of the upper-central target (oscillations from the intended target position to center, further down from center and back) compared to a central one (green).

**Figure 9:**
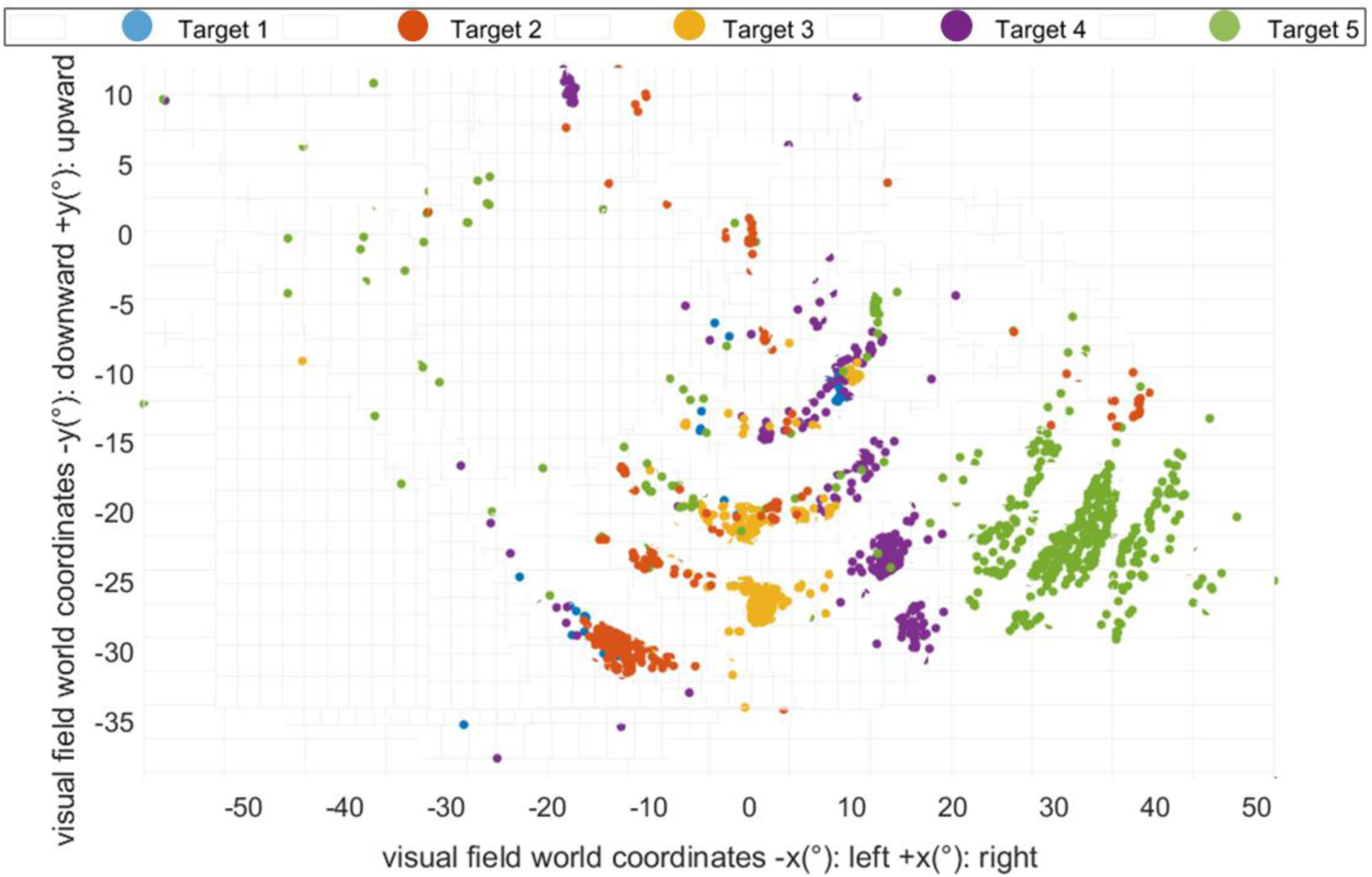
Scatter plot of fixation points of the last row. Starting from left to right target 1 is the most left one positioned in the last row till target 5, the most right one. Blue, orange, yellow, lilac, and green are all fixation points belonging to target 1,2,3,4 and 5 respectively.

Accuracy and precision become worse for different quantiles of users (Table 1). Starting from the 75^th^ quantile both accuracy and precision showed an increase in imprecision, with accuracy passing from a visual angle of 3.21° to 4.88° and 6.06° and precision passing from 1.63° to 2.51° and 3.55° from the 25^th^ quantile to the 75^th^ and 90^th^ quantile respectively.

**Table 1:** Average accuracy and precision across different percentiles of each target in the head-still condition.

|  | Accuracy (°) | Precision (°) |
| --- | --- | --- |
| **Quantile (head-still)** |  |  |
| 25% | 3.21 | 1.63 |
| 50% | 3.98 | 1.95 |
| 75% | 4.88 | 2.51 |
| 90% | 6.06 | 3.55 |

In the head-free condition, there is an overall average of precision of 1.15°, SD: ±0.69. Under head-movement one-way ANOVA revealed a significant difference in precision between Free__stable_, compared to phases of Free__moving_ (F (1,18) = 8.64), p < 0.01; RMS mean__stable_: 0.76°, SD__stable_: ±0.39, RMS mean__moving_: 1.54°, SD__moving_: ±0.74) with a higher imprecision during periods in which subjects were moving their head. As to data loss, there is a double amount of data slippage in the Free__moving_ phase compared to when subjects were not moving their head (7.56% of data spillage compared to 3.69% of data spillage).

### Results: Temporal precision

The one-sample Kolmogorov-Smirnov test showed that the intervals between LED and dot onset (Figure 10) are not extracted from a standard normal distribution, therefore a better indication for comparison between the temporal precision tests is the median (Figure 11).

**Figure 10:**
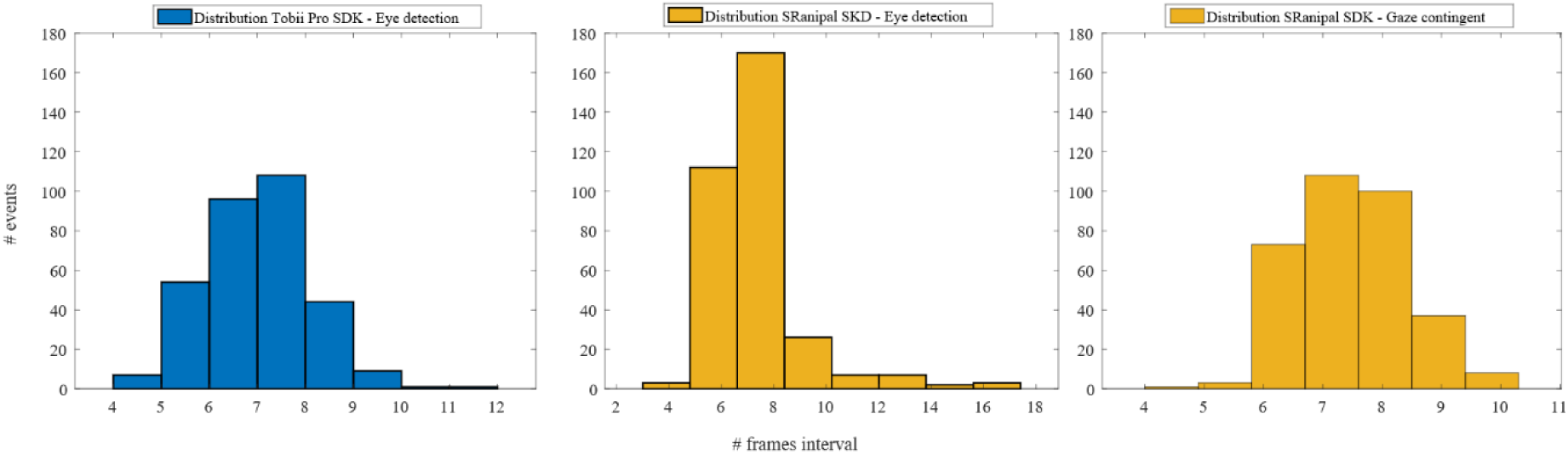
LED-dot frame interval histograms for the eye-detection and the gaze-contingent scenarios. The blue histogram indicates the number of frame distribution when using the Tobii Pro SDK for the LED on–eye on, in the eye-detection scenario (black contour). The yellow histograms show the frame distribution when using the SRanipal SDK. The first yellow one shows the number of frame distribution in the eye-detection scenario (black contour) and the second yellow one in the gaze-contingent scenario (LED on – eye shift).

**Figure 11:**
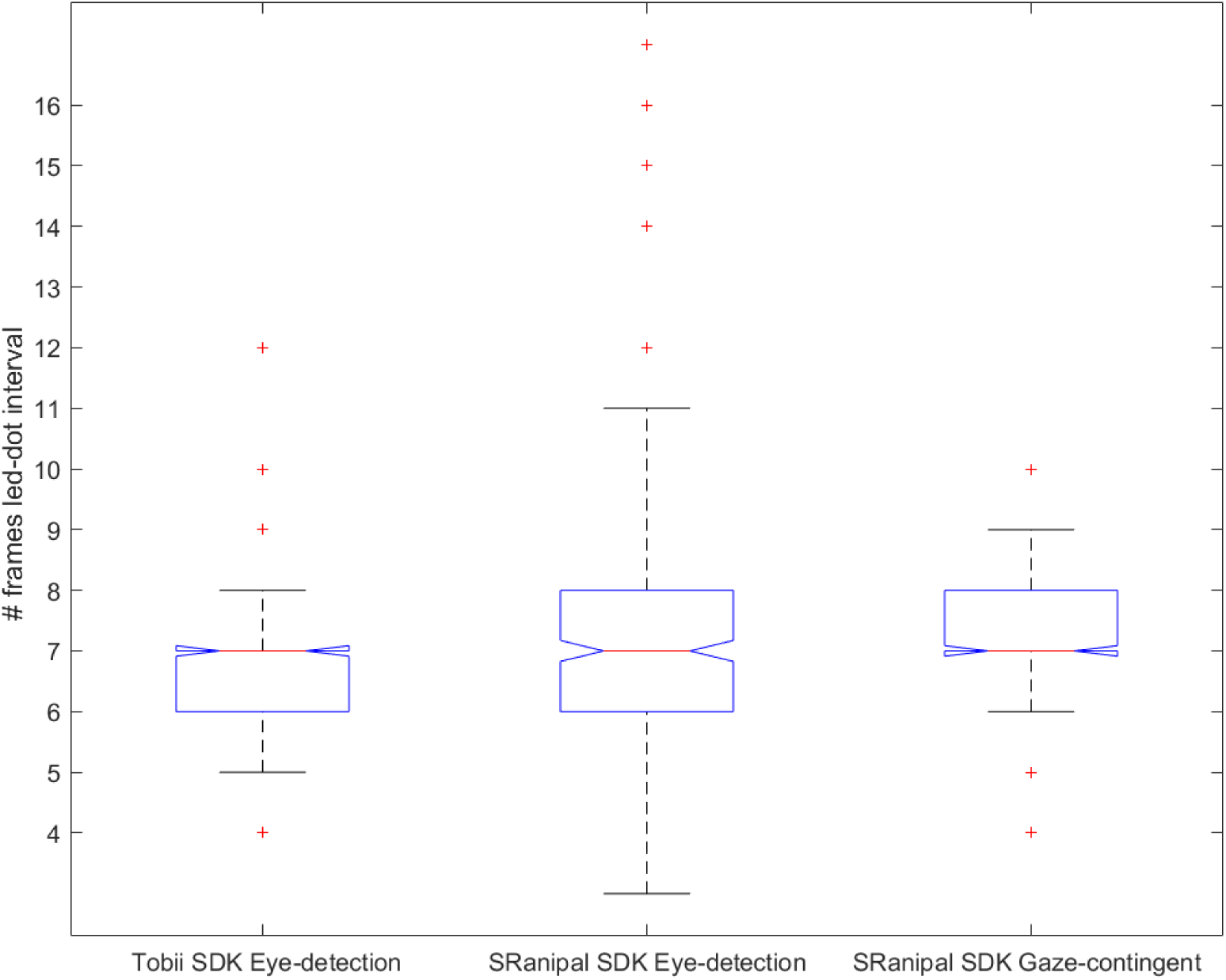
Box plots with LED-dot frame intervals for each scenario. The red line represents the median.

In the eye-detection scenario, for the Tobii Pro SDK and the SRanipal, a median of 58.1 ms is found. In the gaze-contingent scenario, a median temporal precision of 58.1 ms is also found.

## Discussion

We tested data quality and temporal precision of the HTC VIVE Pro Eye in VR to analyze the possible application of the system to VF perimetry testing and simulation of visual loss.

The head-still and head-free conditions showed different limitations of the embedded eye-tracker. The head-still condition evidenced the target position on the screen affected spatial precision. In comparison with the central line, inaccuracy is found for the upper horizontal line and imprecision for the lower one, both around 25° away from the midline. The upper horizontal line shows that fixations in regions above 25° from the midline are difficult. It is hypothesized that subjective facial configurations, such as the distance of the headset from the eyes, is shrinking the visual field and making fixation in that area more challenging. Below 25° from the midline, fixational points are more spread, and the eye-tracker is more imprecise.

In this study, the goal was to examine the precision of a heterogeneous study population. Data quality changes across the population point towards external factors affecting eye-tracking data quality, for example, the used eye correction. Reflections due to eye correction can affect precision when fixating targets placed at the edges (Dahlberg, 2010). Therefore, we hypothesize that the observed deviations could be in part affected by the type of eye correction the subjects were using. To study the influence of the habitual correction was not part of the study, but could be a successor study measuring more subjects.

The head-free condition evidenced how precision and data loss can be influenced by head movement: precision is lower, and a double amount of data loss occurs while moving.

These preliminary results indicate that the status quo of the HTC VIVE Pro Eye has limitations for VR VF perimetry testing. The pilot test evidenced that unfiltered data quality is affected for eccentricities above ±10°, and at ± 25° data is significantly worse. The present results indicate that VR VF testing at ±10° needs improvement. At this eccentricity, the HFA generally is used to detect advanced glaucoma (Asaoka, 2014). Steps to improve accuracy at ±10° are to increase the calibration points and position them at of areas of interest for the test (Holmqvist et al., 2012). The strong limitations observed at ± 25° indicate that the HTC Vive Pro Eye for VR perimetry testing to detect the early onset of glaucoma (Nouri-Mahdavi, 2014) is very restricted.

As to passive contingent studies that specifically require a small temporal precision, preliminary conditions should be kept in mind. An acceptable level of system’s latency depends on the application. Ideally, the display should be updated immediately at the end of each saccade. In practice, this is limited since a lag always exists between identification of saccade ending, rendering the new image, transmitting it, and displaying it (Loschky and Wolverton, 2007). For example, rendering the image can take from 25 up until 150 ms (Thomas and Geltmacher, 1993; Ohshima, et al., 1996; Geisler and Perry, 1998).

Furthermore, the display refresh rate can make a difference between a good or an acceptable level of latency (Saunders and Woods, 2014; Gibaldi et al., 2017). The eye tracker used in the HTC Vive Pro Eye has a higher refresh rate than the display, therefore, for this system, one part of the latency’s variance can be due to the display’s refresh. The direct, objective and automatic temporal precision tests showed that there is no difference between the detection of an eye and a gaze-contingent scenario. More so, displaying data through the Tobii Pro SDK or the SRanipal SDK makes no difference in terms of temporal precision.

For all the tests conducted, the median is a good indicator of temporal precision. The value of 58.1 ms makes the system suitable for one type of passive gaze-contingent application: multiresolution displays that can be applied to simulate tunnel vision (Stock et al., 2019). Multiresolution displays are generally used for foveated rendering, and latencies between 50 and 70 ms are well accepted because modifications are made in the periphery, and they are not usually detected (Loschky and Wolverton, 2007; Albert et al., 2017). The reason why that happens is that changes in the postsaccade area mostly overlap with the changes in the presaccadic one (Saunders and Woods, 2014). On the other hand, if changes are made in the central area, such as for VR studies simulating scotomas (Ai et al., 2000; Banks and McCrindle, 2008; Lewis et al., 2012; Jin et al., 2016; Väyrynen et al., 2016; Wu et al., 2018), where a mask is applied to central vision, a smaller latency is required. The postsaccade target is not already masked, therefore with latencies longer than saccade durations, there is the danger of central-vision hint of the target (Saunders and Woods, 2014).

For these applications, the HTC Vive Pro Eye needs improvement. Steps to improve latency in scotoma-simulated gaze-contingent VR studies are to predict where the saccade will likely end instead of using the current gaze position (Arabadzhiyska et al., 2017), or to apply bigger scotoma sizes. In the latter case, the reduction in latency is dependent upon saccades distances concerning the scotoma size (Saunders and Woods, 2014).

## Conclusion

The present preliminary study shows that thanks to a temporal precision of 58.1 ms, HTC Vive Pro Eye is suited for peripheral visual field loss simulation and needs improvements for central visual loss simulation. The system needs adjustments also for advanced glaucoma VR VF perimetry detection and shows high limitations for early glaucoma detection onset due to eye-tracking data inaccuracy and impression at the custom visual field testing used by the HFA.

## Acknowledgments

This work was funded by the Federal Ministry of Education and Research of Germany in the framework of IDeA (project number 16SV8104). The authors acknowledge intra-mural funding of the University of Tübingen through the mini graduate school ‘Integrative Augmented Reality (I-AR)’.

